# Soft tissue deformations explain most of the mechanical work variations of human walking

**DOI:** 10.1101/2021.04.14.439866

**Authors:** Tim J. van der Zee, Arthur D. Kuo

## Abstract

Humans perform mechanical work during walking, some by leg joints actuated by muscles, and some by passive, dissipative soft tissues. Dissipative losses must be restored by active muscle work, potentially in amounts sufficient to cost substantial metabolic energy. The most dissipative, and therefore costly, walking conditions might be predictable from the pendulum-like dynamics of the legs. If pendulum behavior is systematic, it may also predict the work distribution between active joints and passive soft tissues. We therefore tested whether the overall negative work of walking, and the fraction due to soft tissue dissipation, are both predictable by a pendulum model across a wide range of conditions. The model predicts whole-body negative work from the leading leg’s impact with ground (termed the Collision), to increase with the squared product of walking speed and step length. We experimentally tested this in humans (N = 9) walking in 26 different combinations of speed (0.7 – 2.0 m·s^-1^) and step length (0.5 – 1.1 m), with recorded motions and ground reaction forces. Whole-body negative Collision work increased as predicted (R^2^ = 0.73), with a consistent fraction of about 63% (R^2^ = 0.88) due to soft tissues. Soft tissue dissipation consistently accounted for about 56% of the variation in total whole-body negative work. During typical walking, active work to restore dissipative losses could account for 31% of the net metabolic cost. Soft tissue dissipation, not included in most biomechanical studies, explains most of the variation in negative work of walking, and could account for a substantial fraction of the metabolic cost.

**Summary statement:** Soft tissue deformations dissipate substantial energy during human walking, as predicted by a simple walking model.

## Introduction

Human walking costs metabolic energy in part because the muscles perform positive work to counter the negative work within each stride. There is currently no mechanistic prediction for how much work is performed, except that the positive and negative work of a steady stride cancel each other, so that the negative work could be considered predictive of the positive work. Some of the negative work occurs actively when muscles act eccentrically to lengthen under load, and some occurs as passive dissipation from soft tissue deformations (Fig. 1A). Passive dissipation may account for 31% of the negative work of typical walking (Zelik & Kuo, 2010), but its distribution relative to active negative work is not known for more general walking conditions. However, walking patterns have been observed to scale quite consistently across a wide range of gait parameters (Grieve, 1968), and total negative work to be predictable (Adamczyk & Kuo, 2009). This suggests that the amount of passive dissipation and active negative work may be predictable across gait conditions. Such predictability could provide insight on when muscles must actively perform positive work and thus consume metabolic energy.

**Fig. 1:**
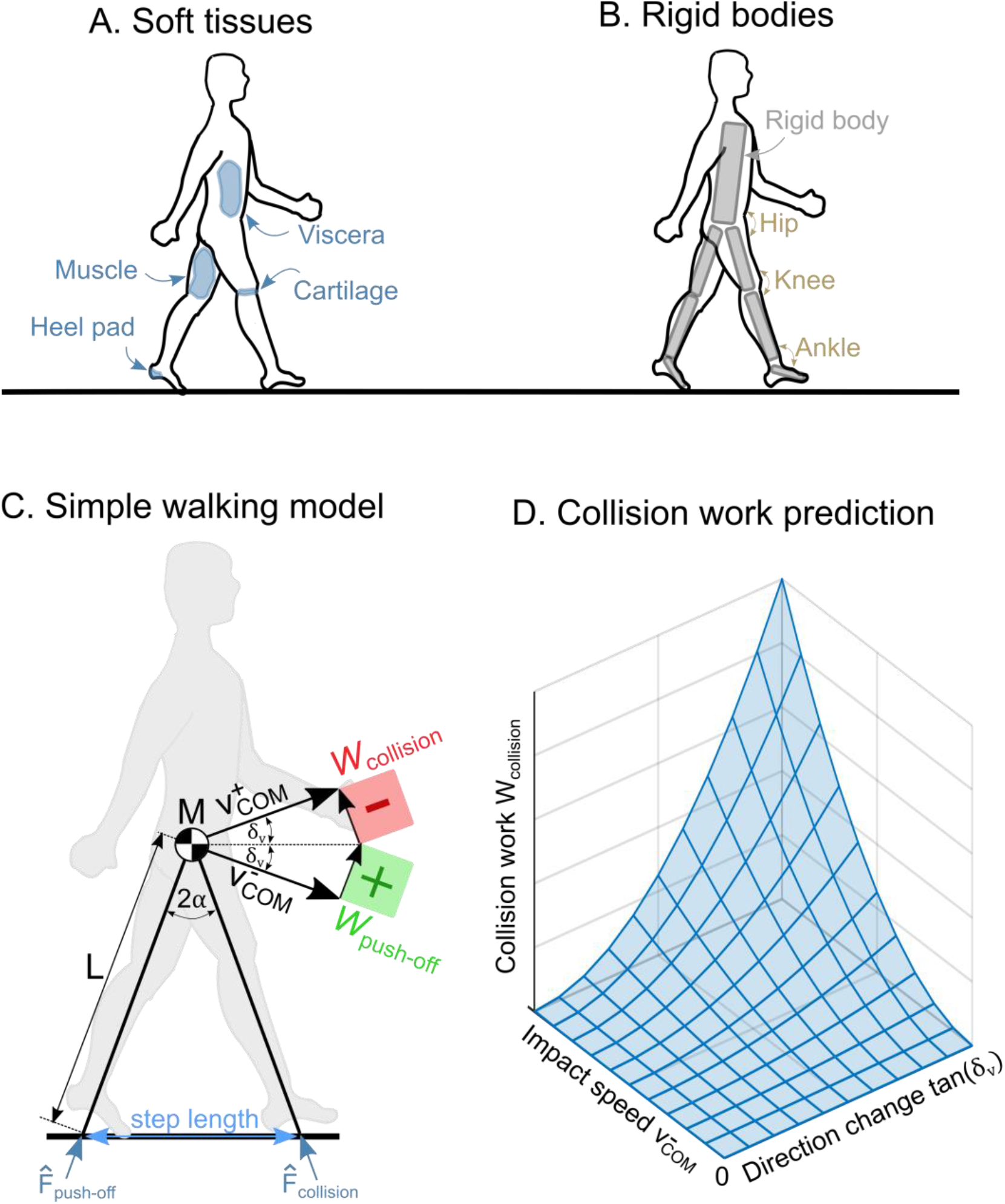
Soft tissues, rigid bodies and negative work predicted by simple walking model. **A: Soft tissues** such as viscera, muscle, cartilage and heel pad can dissipate energy by delivering force while deforming and/or displacing. **B: Rigid bodies** are traditionally used for estimating joint torques and work (rates), using an approach referred to as inverse dynamics. **C: Simple walking model** predicts negative Collision work *W*_collision_ at heel strike from center of mass (COM) velocity *v*_COM_. A portion of this negative work may be due to soft tissue dissipation. **D: Predicted Collision work** *W*_collision_ increases with impact speed 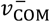 squared and the tangent of COM velocity directional change tan(*δ*_*v*_) squared.

One of the critical events of walking is the impact of the leg with ground after the swing phase. After heel strike, the leading leg performs negative work (during a phase termed Collision) as the body center of mass (COM) velocity is redirected from a forward-and-downward direction from the previous stance phase, to a forward-and-upward direction at the beginning of the next (Adamczyk & Kuo, 2009; Kuo et al., 2005). For typical walking at 1.25 m · s^−1^, about 12.5 J of negative work is done during Collision within the first 15% of the stride, with contributions from active muscle-tendons and passive soft tissues (about 40% and 60%, respectively; Zelik & Kuo, 2010). The soft tissues responsible for the dissipation are thought to include the foot and shoe (Honert & Zelik, 2019), particularly the heel pad (about 3.8 J; Baines et al., 2018), as well as the shank (Pain & Challis, 2001). Some passive dissipation may also occur with loading of articular cartilage (Hayes & Mockros, 1971) and intervertebral discs (Virgin, 1951), and from inertial loading of wobbling mass, for example muscle (Schmitt & Günther, 2011) and viscera (Minetti & Belli, 1994). Soft tissue dissipation appears to vary consistently with overall collision work, for example in obese and non-obese adults (Fu et al., 2015), and even for landing from a jump (Zelik & Kuo, 2012). The overall Collision work accounts for most of the negative (and thereby positive) work of a stride (Zelik & Kuo, 2010), but it is unknown how it varies with gait conditions, and particularly how much of it is due to soft tissue dissipation.

The remainder of the stride appears to be systematically related to Collision. Following Collision negative work are alternating phases of positive and negative work by the whole body. The work done during these phases (termed Rebound, Pre-load, and Push-off during stance (Donelan et al., 2002; Zelik & Kuo, 2010) increases in proportion to Collision during walking at preferred step length (Zelik & Kuo, 2010). These alternating phases resemble the oscillation of a purely elastic spring for each leg (Geyer et al., 2006), excited by ground contact. In reality, that action is performed not by springs, but by a combination of active muscle-tendons and passive soft tissues. The oscillatory behavior suggests that muscles are actively controlled as a function of dynamical state, so that the entire body acts like a consistent dynamical system. The negative work of an entire stride might therefore vary systematically with the magnitude of Collision work, and across a variety of gait conditions.

The amount of Collision work, and by extension of the entire stride, may actually be predictable (Fig. 1D). A simple dynamic walking model predicts how work must be performed on the body center of mass (COM) to redirect its velocity between pendulum-like stance phases (Kuo, 2001), forward-and- downward at the end of one arc, to forward-and-upward at the beginning of the next. This is achieved by negative Collision work with the leading leg, and positive push-off work from the trailing leg. The Collision work varies with gait parameters such as walking speed, step length, and step frequency (Adamczyk & Kuo, 2009), as predicted by the simple model (Kuo, 2001). The work of the other phases may vary in proportion to the Collision work, if part of a dynamical system. Similarly, the active and passive work may also vary in proportion to the Collision work. The soft tissue dissipation during Collision and the total negative work over a full stride might therefore be proportional to predictions from the simple walking model.

The purpose of the present study was to test whether the negative work of walking is systematically distributed between active and passive contributions, and across the entire stride. The starting point for this inquiry is the whole-body Collision work, as previously modeled and characterized across gait parameters such as step length and step frequency (Adamczyk & Kuo, 2009). For active vs. passive contributions, we hypothesized that (passive) soft tissue dissipation during Collision would remain a consistent fraction of the whole-body Collision work predicted by model. And for the entire stride, we hypothesized that whole-body Collision work would remain a consistent fraction of the negative work over a full stride. We tested these predictions with a human walking experiment, in which whole-body and soft tissue work were estimated for 26 different combinations of gait parameters, including a range of walking speeds while constraining step length and/or step frequency. The results may indicate whether simple pendulum-like walking models can predict both the amount and distribution of negative work in humans.

## Methods

We used a simple walking model to predict the Collision work during walking, and tested whether it was predictive of soft tissue dissipation during Collision, as well as to overall negative work for the full stride. We tested predictions against experimental data on human subjects walking across a wide range of combinations of walking speed and step length. The testing data included rigid body mechanical work rate from inverse dynamics (see Fig. 1B), as well as whole-body and soft tissue mechanical work rates from a previously-reported procedure based on COM work and motion capture (Zelik & Kuo, 2010, 2012). We next present the model predictions, followed by the experimental tests.

### Model Predictions

Predictions for negative Collision work *W*_collision_ were produced from a simple walking model (Kuo, 2001; Adamczyk & Kuo, 2009), see Fig. 1C. The stance leg is treated as a simple inverted pendulum with length *L* and body mass M concentrated at the pelvis, and the swing leg as a simple pendulum with length *L* and infinitesimal mass foot. When the swing leg hits the ground, a Collision impulse 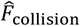 does negative Collision work *W*_collision_ on the COM,

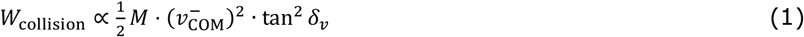

where *M* is body mass, 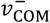 pre-impact COM speed, and *δ*_*v*_ the COM velocity directional change (Fig. 1C). The Collision work *W*_collision_ in humans is hypothesized to increase similar to the simple model’s, with an empirical proportionality due to unmodeled effects such as imperfectly rigid legs, distributed body mass, and a finite-length (as opposed to point) foot (Adamczyk et al., 2006). The Collision work may thus be regarded as a function of impact speed 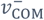 and COM velocity directional change *δ*_*v*_ (Fig. 1D).

We hypothesize that the mechanical work of walking is distributed systematically between active joints and passive soft tissues, and across the time of a stride. Both soft tissues and joints (actuated by active muscles in series with tendons) contribute to total Collision work. If the distribution is systematic, the soft tissue negative work during Collision *W*_soft,collision_ will be responsible for a consistent fraction of the Collision work W_collision_:

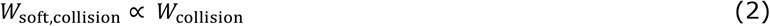

Systematic distribution also means work should be distributed proportionately within a stride. This means that total negative work over the entire stride *W*_stride_ will change in proportion to the Collision work *W*_collision_:

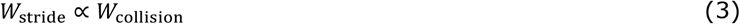

These hypotheses lead to several expectations for human experiments. For Collision work *W*_collision_, we introduce an empirical coefficient *c*_collision_ for the proportionality (Eq. 1)

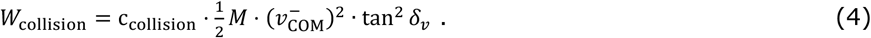

For soft tissue Collision work *W*_soft,collision_, its proportionality (Eq. 2) to total Collision work *W*_collision_ results in the expectation

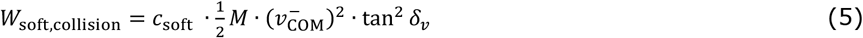

with coefficient *c*_soft_. There may also be work within a full stride not related to Collision (Eq. 3). For example, the knee performs negative work to decelerate the swing leg, which has little effect on the COM. Such contributions are expected to have little or no dependency on the COM velocity, and are therefore lumped into a single constant offset *d*_stride_:

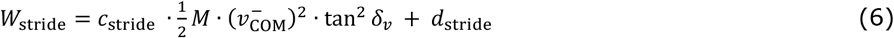

Altogether, we expect that soft tissues contribute to Collision work *W*_collision_, with a remainder explained by active joints. We expect that Collision work *W*_collision_ contributes to full stride negative work *W*_stride_, with a remainder due to negative work during pre-load and swing phases. We therefore expect soft tissue Collision work *W*_soft,collision_ to be smaller than Collision work *W*_collision_, which we expect to be smaller than full stride negative work *W*_stride_, such that *c*_stride_ > *c*_collision_ > *c*_soft_. We test for *c*_stride_, *c*_collision_ and *c*_soft_ using regression on experimental data. As this set of predictions depends entirely on velocity (i.e.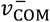 and *δ*_*v*_), we refer to it as velocity-based predictions.

We also tested another set of gait-based predictions rather than velocity data. Gait parameters such as average speed 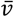 and step length *s* are usually more readily available than velocity data, and can also serve as predictors. This requires an additional set of assumptions, that average speed 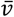 and step length *s* are respectively proportional to impact speed 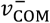 and the tangent of COM velocity directional change tan *δ*_*v*_. With a small angle approximation, step length is proportional to the inter-leg angle *α* (Fig. 1C), which should equal *δ*_*v*_, and with another small angle approximation, tan *δ*_*v*_. Thus, all gait-based predictions are

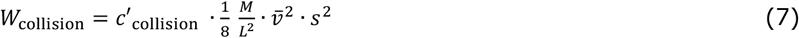

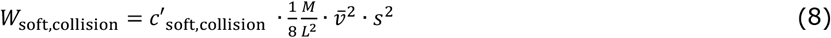

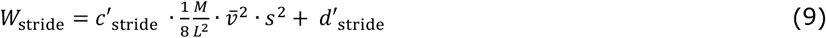

where the prime symbol (′) refers to gait-based predictors. As with the original coefficients, we expect that *c*′_stride_ > *c*′_collision_ > *c*′_soft_, tested using regression on experimental data.

### Experimental Procedures

Healthy adult subjects (*N* = 9, body mass *M* 73.5±15 kg, leg length *L* 0.93±0.06 m, age 23.5±2.5 years, mean ± s.d.) walked on an instrumented treadmill at 26 different combinations of walking speed and step length. The combinations belonged to four experimental conditions: *Preferred walking* at varying walking speeds 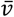, *Fixed frequency* at varying step lengths *s, Fixed step length* at varying frequencies *f*, or *Fixed average speed* with inversely varying combinations of step length and step frequency (see Table 1). Step length *s* and step frequency *f* were varied relative to the preferred values *s*\* and *f*\*, determined from unconstrained walking at a nominal speed (*v*\* = 1.25 m/s). Walking speed 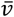 and step frequency *f* were manipulated by setting the treadmill belt speed and asking subjects to walk on the beat of an audio cue, respectively. Step length *s* was manipulated through both walking speed and step frequency from their ratio 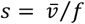. The order of trials was randomized for each subject individually, who were earlier familiarized with each of the conditions during a 6-minute practice trial. Subjects provided their informed consent to participate in the experiment, which was approved by the Institutional Review Board of the University of Michigan, where the experiment was performed.

**Table 1:**
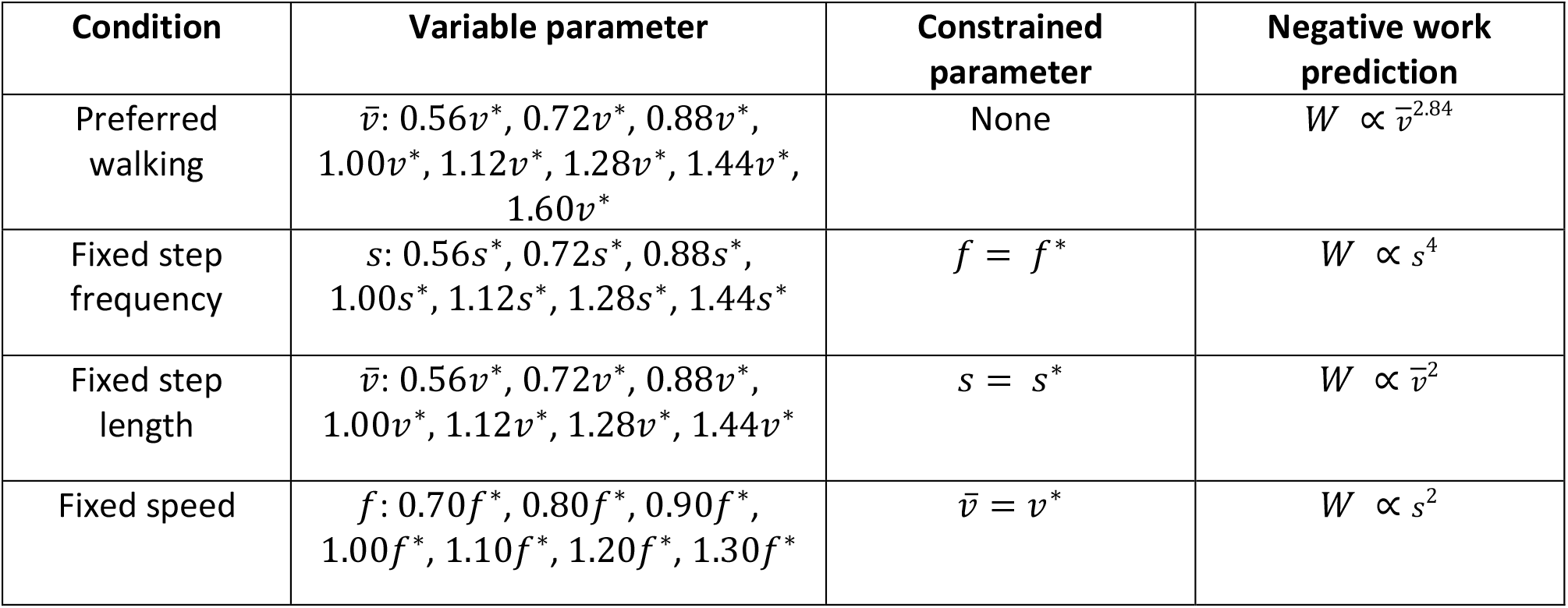
Each of four experimental conditions including preferred walking with no constraint, and others constraining one of frequency, step length or walking speed.

We also used the gait-based coefficients *c*′_stride_, *c*′_collision_ and *c*′_soft_ to evaluate condition-specific predictions. For preferred and fixed step length walking, step length *s* is expected to increase with 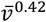 (Grieve, 1968) and 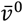 respectively so that predicted dissipation increases with 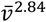 and 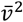 respectively (see Fig. 2 and Table 1). For fixed step frequency and fixed speed walking, average speed 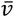 increases with *s*^1^ and *s*^0^ respectively so that predicted dissipation increases with *s*^4^ and *s*^2^ respectively.

**Fig. 2:**
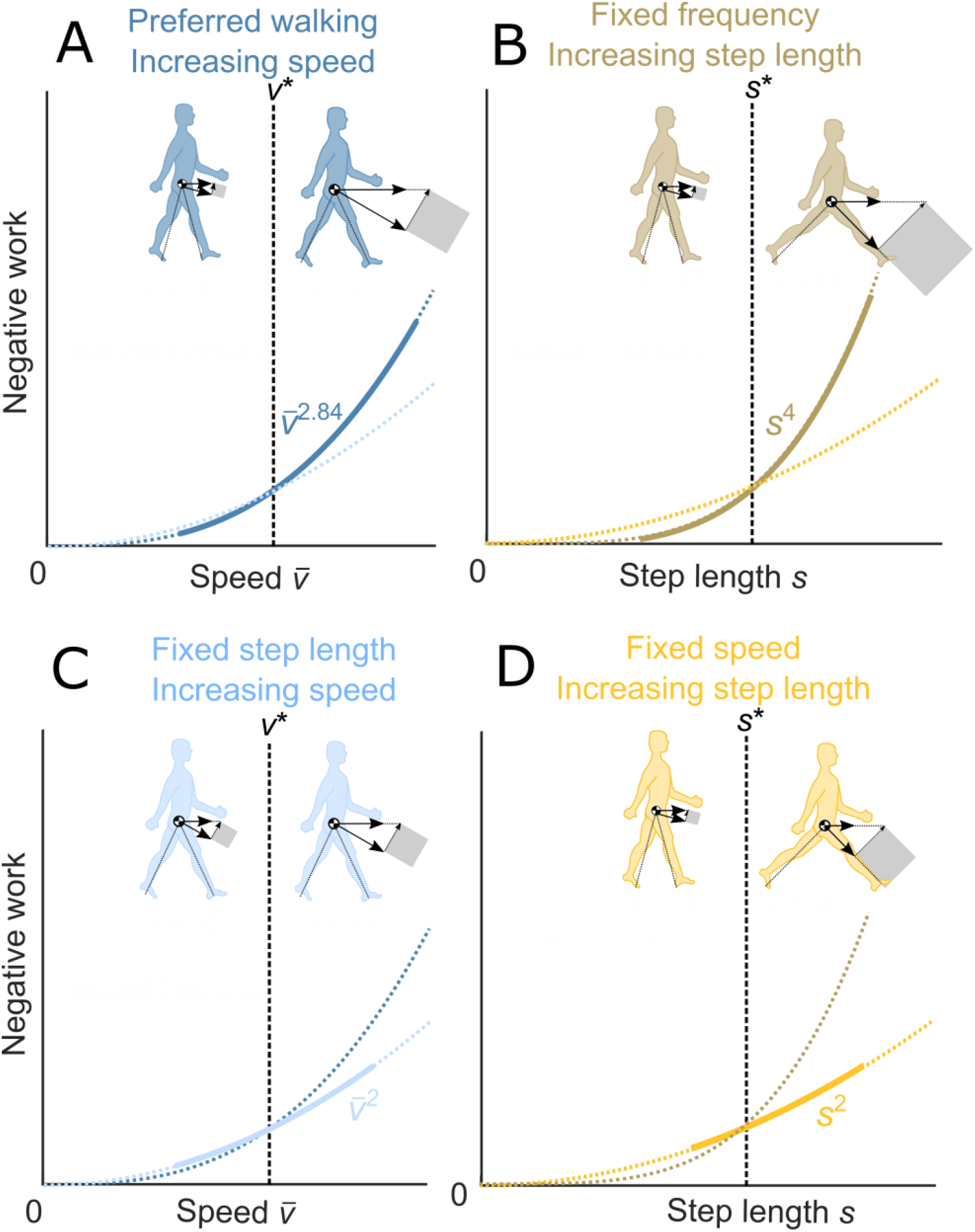
Predictions of negative work during walking. **A. Preferred walking:** Assuming the empirical preferred step length relationship, negative work increases with (speed)^2.84^. **B. Fixed frequency:** If speed increases linearly with step length, negative work increases with (step length)^4^. **C. Fixed step length:** If step frequency increases linearly with step frequency, negative work increases with (speed)^2^. **D. Fixed speed:** If step length and frequency vary inversely at fixed speed, negative work increases with (step length)^2^. These relationships are predicted to apply to total Collision work, soft tissue Collision work, and total negative work over a full stride.

### Human experiment

We used ground reaction forces and motion capture to estimate work performed by the body, including rigid body and soft tissue work. Ground reaction force *F*_gr_ was measured from the instrumented treadmill (Bertec) at 1200 Hz and low-pass filtered at 25 Hz. Instantaneous COM velocity *v*_COM_ was determined from integrating the ground reaction force *F*_gr_ using a periodicity assumption. Center of mass work rate 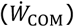 was determined from the dot product of each leg’s ground reaction force *F*_gr_ and center of mass velocity *v*_COM_ (Donelan et al., 2002). In addition to ground reaction force measurements, motions of the lower limbs were captured using Motion Analysis at 120 Hz. Cluster markers were attached to the feet, shanks and thighs. Single markers were located at the head of the 5^th^ metatarsus, left and right malleoli, left and right epicondyles, greater trochanter, left and right anterior superior iliac spine and sacrum. Ankle, knee and hip joints were defined based on locations of malleoli, epicondyles and Helen-Hayes Davis points respectively (Davis et al., 1991). Ground reaction force and motion capture measurements were used in standard inverse dynamic analysis (Visual 3D, C-Motion, Germantown, MD, USA) for computing the joint angles, translational displacements, torques, forces and work rates in the ankle, knee and hip. The joint work rates were summed across joints to yield overall rigid body work rate 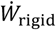 (also referred to as Summed Joint Power using six-degree of freedom joints, (e.g. Honert & Zelik, 2019)). In addition, the segmental kinetic energies of the feet, shanks and thighs were computed assuming rigid bodies. At least five strides from each condition were analyzed per participant, selected to avoid motion capture occlusions and steps that landed on both force plates. Some of the trials could not be analyzed (five out of 324) due to missing data (two), incorrect stepping (one) or lose markers (two). These occasions all belonged to different subjects, and to different conditions. An additional (tenth) subject was recorded in experiment but excluded from analysis owing to incorrect pelvis marker placement.

The resulting data were then used to estimate work quantities, as described previously (Zelik et al., 2015; Zelik & Kuo, 2010). Whole-body work rate 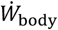 was defined as the COM work rate 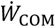, plus the peripheral work rate 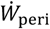, defined as the sum of all unilateral segmental kinetic energy fluctuations about the COM. The trunk and upper limbs contribute relatively little to walking (Vaughan et al., 1992), and so we limited our segmental analysis to the lower limbs (Fu et al., 2015). Whole-body work rate 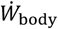 typically becomes negative during the *Collision* phase right after heel strike, then becomes positive during *Rebound* until mid-stance, when it becomes negative during *Pre-load*, before a final positive-work *Push-off* at end of stance (Donelan et al., 2002). The phases other than Collision are mainly used for qualitative illustrative purposes, although Pre-load and Collision together contribute to the overall negative work of a stride, *W*_stride_. Collision phase was defined as the interval between heel-strike and the instant of the steepest upward COM velocity (Adamczyk & Kuo, 2009). Soft tissue work rate 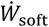 was determined from the discrepancy between rigid body work rate 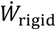 and whole-body work rate 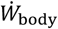 (Zelik et al., 2015; Zelik & Kuo, 2010). All work rates were calculated for each leg individually and then averaged between legs.

The primary work quantities of interest were Collision work *W*_collision_, soft tissue Collision work *W*_soft,collision_, and total negative work over the entire stride *W*_stride_. Collision work *W*_collision_ and soft tissue Collision work *W*_soft,collision_ were obtained by time-integrating whole-body work rate 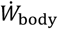 and soft tissue work rate 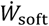 respectively, for the negative intervals of 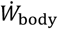 during Collision. Total negative work per stride *W*_stride_ was obtained from time-integrating whole-body work rate 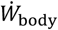 during the intervals when it was negative. All of these work quantities were negative and reported as magnitudes and computed as work per stride to facilitate comparison to the simple walking model.

## Results

The measured quantities generally exhibited qualitatively systematic variations with the gait conditions. For example, sagittal plane joint angle, torque and rotational work rate during walking at constant step frequency and increasing step length generally increased in amplitude with walking speed (see Fig. 3). Each joint’s contribution may be inferred by comparing its work rate to the rigid body work rate (see Fig. 3). Each joint contributed differently to rigid body work rate, with mainly positive contributions from the ankle, negative contributions from the knee and mixed contributions from the hip. Whole-body and rigid body work rates during preferred walking at 1.25 m s^-1^ were similar in shape, but different in amplitude. These work rates and their difference were used to determine soft tissue Collision work (see Fig. 4).

**Fig. 3:**
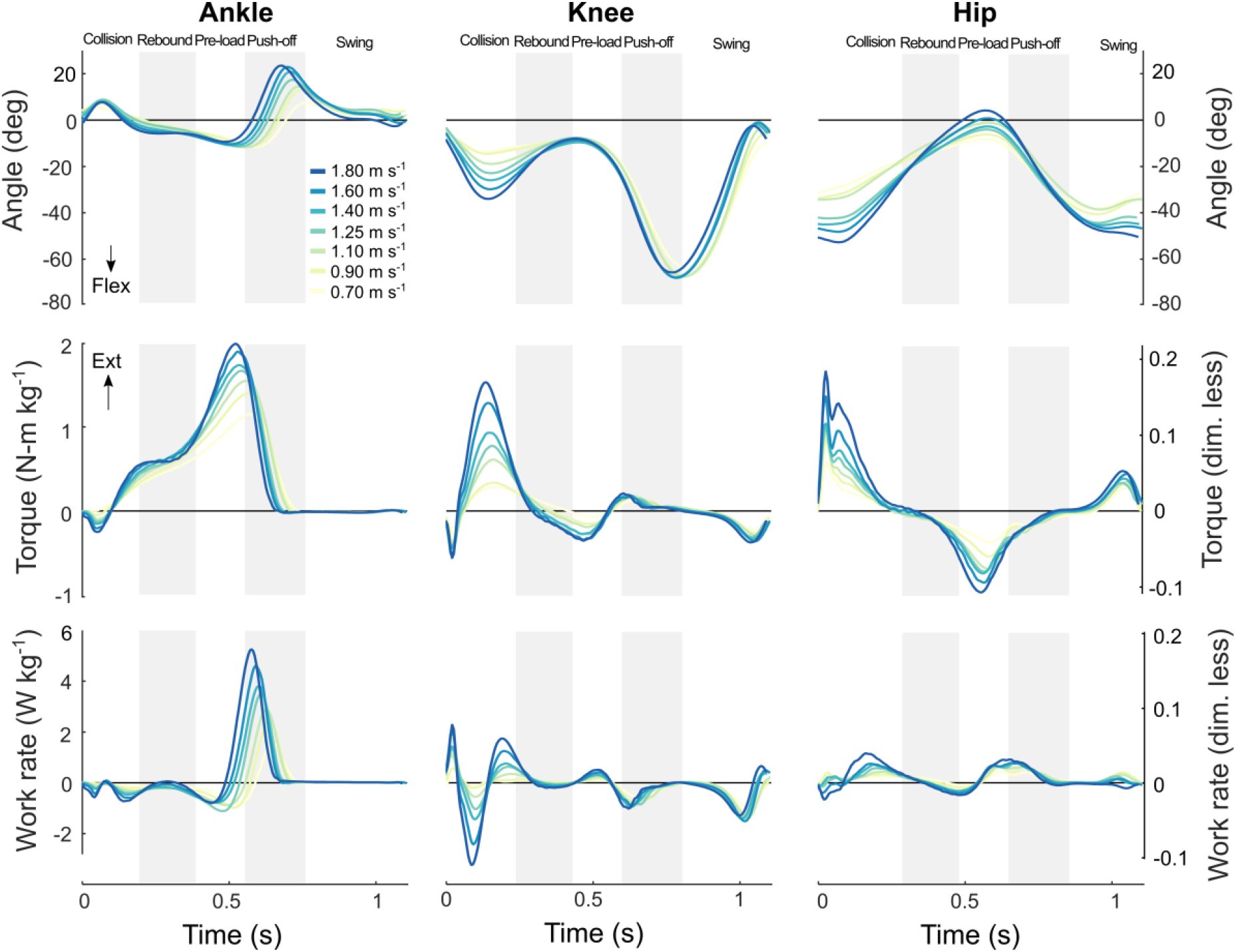
Joint angle, torque, and work rates during walking at constant step frequency. Amplitudes of sagittal plane joint angle, joint torque and joint work rate increased with walking speed. Increases were most pronounced during collision (knee and hip) and push-off phases (ankle and hip).

**Fig. 4:**
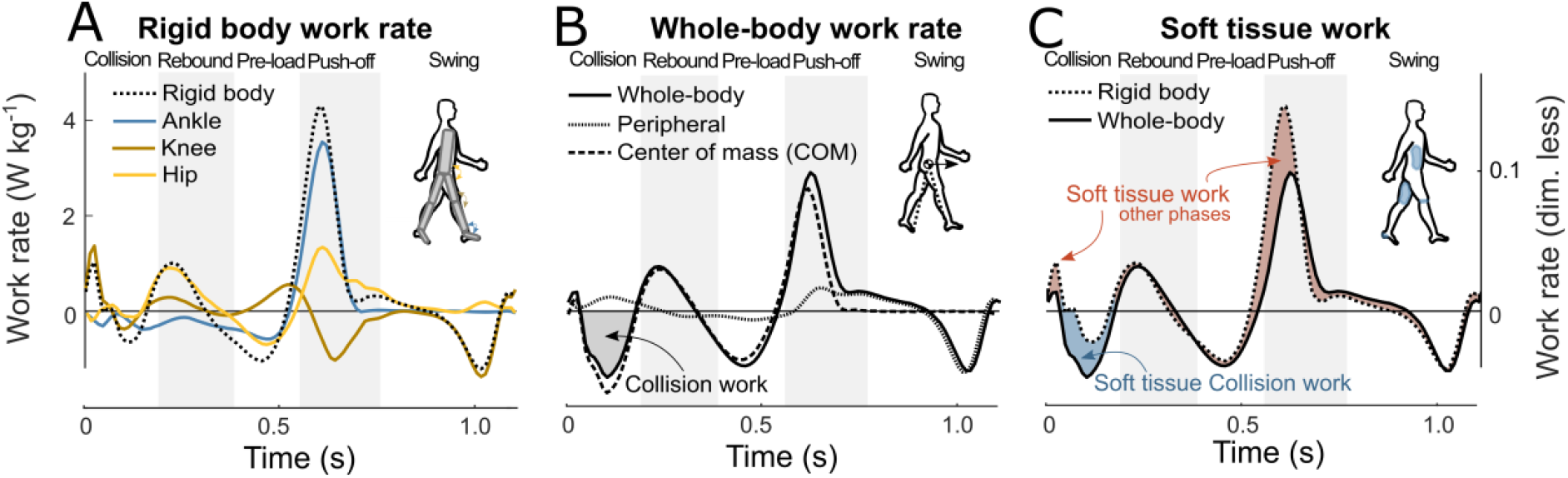
Work rate vs. time for three measures: **(A.) Rigid body, (B.) Whole-body, and (C.) Soft tissue.** Shown are representative data for preferred walking at 1.25 m s^-1^. (A.) Inverse dynamics yields rigid body work rate (dotted line), defined as the sum of work rates from ankle, knee and hip (solid, colored lines). (B.) Whole-body work rate is the sum of center of mass (COM) work rate and peripheral work rate (Zelik & Kuo, 2010). Collision work (shaded region) is the negative whole-body work during Collision. (C.) Soft tissue work rate is the difference between whole-body work rate and rigid body work rate, also with soft tissue work quantified (shaded regions).

Net rigid body work was generally positive and appeared to increase with speed and step length (see Fig. 5). This is consistent with the expectation that soft tissues perform net dissipative work, not captured by rigid body inverse dynamics. In preferred walking and fixed step length walking, net rigid body work per stride appeared to increase with walking speed, with positive contributions from ankle and hip and negative contributions from knee (Fig 5A and 5C). The effect of speed on hip and knee joint work was more pronounced in fixed step length walking (Fig 5C), whereas the effect on ankle joint work was more pronounced in preferred walking (Fig 5A). In fixed frequency walking, net rigid body work per stride appeared to increase with step length, with positive contribution from ankle and nearly constant contributions from hip (positive) and knee (negative) (Fig 5B). In fixed speed walking, net rigid body work per stride appeared to increase with step length, with mainly positive contributions from ankle and mixed contributions from knee and hip. The latter two work terms showed a minimum and maximum at intermediate step lengths respectively, increasing in amplitude towards the extremes.

**Fig. 5:**
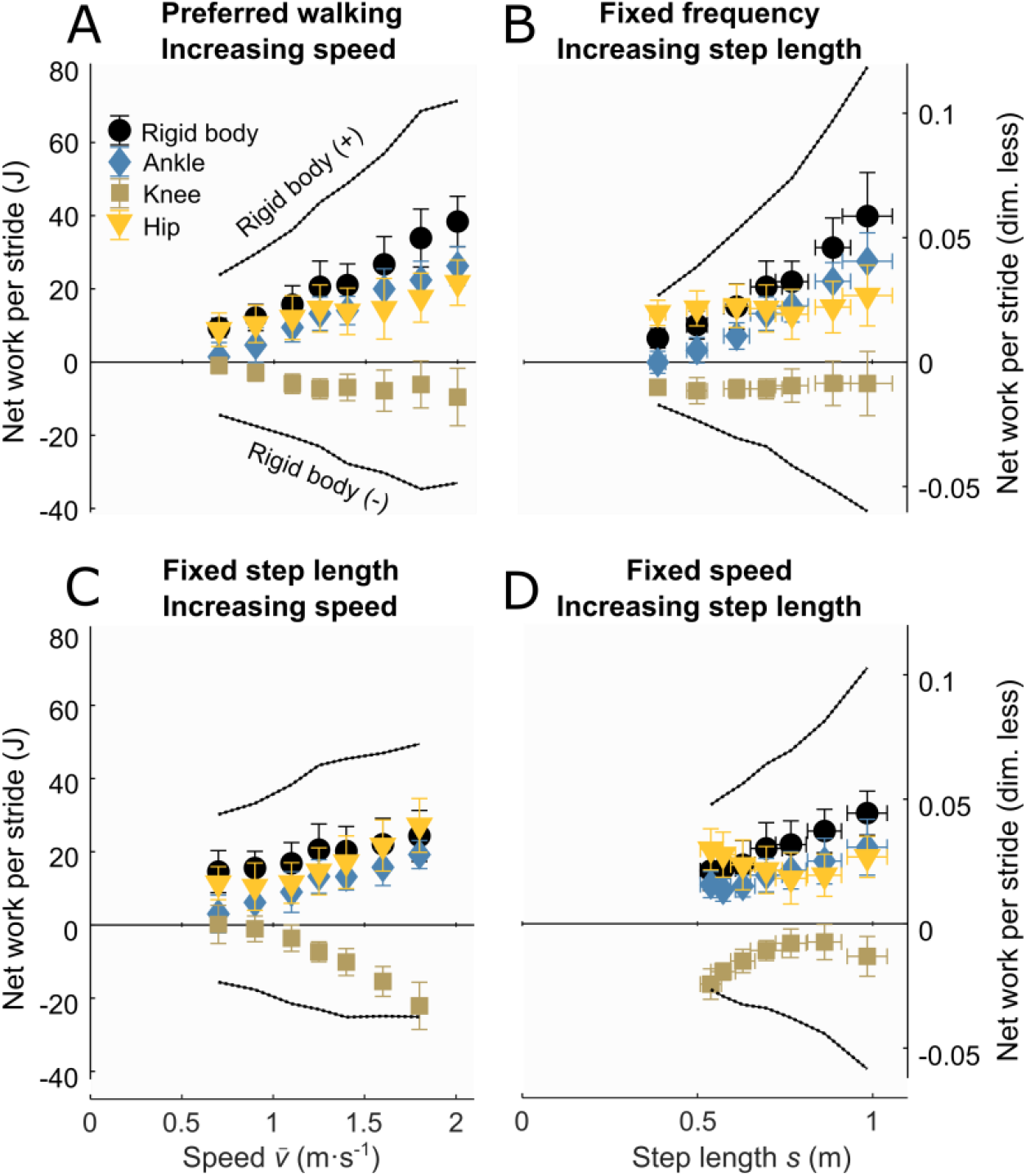
Net joint work per stride during walking in four different conditions. A. Preferred walking at increasing speed. B. Fixed frequency with increasing step length and speed C. Fixed step length with increasing speed and step frequency. D. Fixed speed with increasing step length and decreasing step frequency. Shown are Rigid body work, which is the sum of Ankle, Knee, and Hip joint work per stride. Envelopes (dashed lines) indicate Rigid body positive (+) and negative (–) work, defined as sum of all positive (negative) work from each joint.

Whereas all work rate amplitudes generally increased with speed and step length, the increase was more pronounced for whole-body work rate than for rigid body work rate (see Fig. 6-7). This increase was most notable during collision and push-off phases. Rigid body work rate was similar in shape as whole-body work rate, but different in amplitude (upper panels versus middle panels Fig. 6-7). During collision phase specifically, whole-body work rate had a larger amplitude than rigid body work rate, resulting in a substantial soft tissue work rate amplitude (lower Fig. 6-7). This soft tissue work rate amplitude seemed to increase with step length (lower panels Fig. 6) and with walking speed (lower panels Fig. 6-7), similarly as the amplitude of the whole-body work rate (upper panels Fig. 6-7). Altogether, whole-body and soft tissue work rate amplitudes during collision phase seemed to increase with walking speed and step length to a larger extent than rigid body work rate amplitudes. Soft tissue and rigid body work rates differed in the response immediately after Collision. In nearly all conditions, soft tissue work rate returned to nearly zero at the end of Collision, with little or no positive work. In contrast, the rigid body work typically became positive after Collision, during the Rebound phase. We therefore interpret the soft tissue Collision work as being largely passive and dissipative, whereas the rigid body Collision work may have both active and passive contributions, including a possible damped elastic oscillation. In the remaining, soft tissue Collision work is therefore also referred to as soft tissue dissipation.

**Fig. 6:**
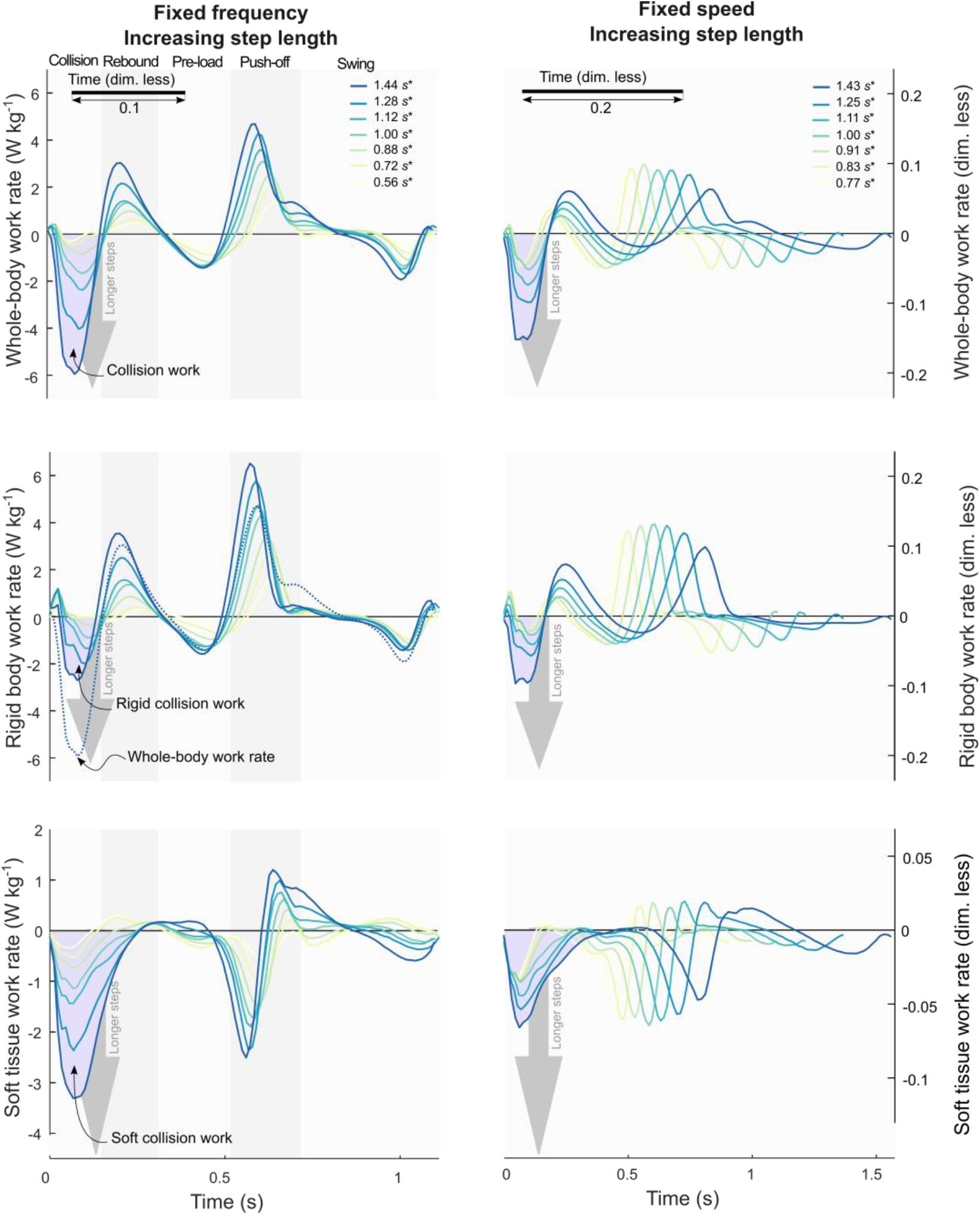
Work rates for walking with (left:) Fixed step frequency and (right:) Fixed speed, both with increasing step length. (top to bottom:) Whole-body, Rigid body, and Soft tissue work rates vs. time for one stride, with shaded regions indicating Collision phase. Whole-body work rate is sum of COM and peripheral work rates. Rigid body work is sum of joint work rates from inverse dynamics. Soft tissue work rate is difference between Whole- and rigid body rates.

**Fig. 7:**
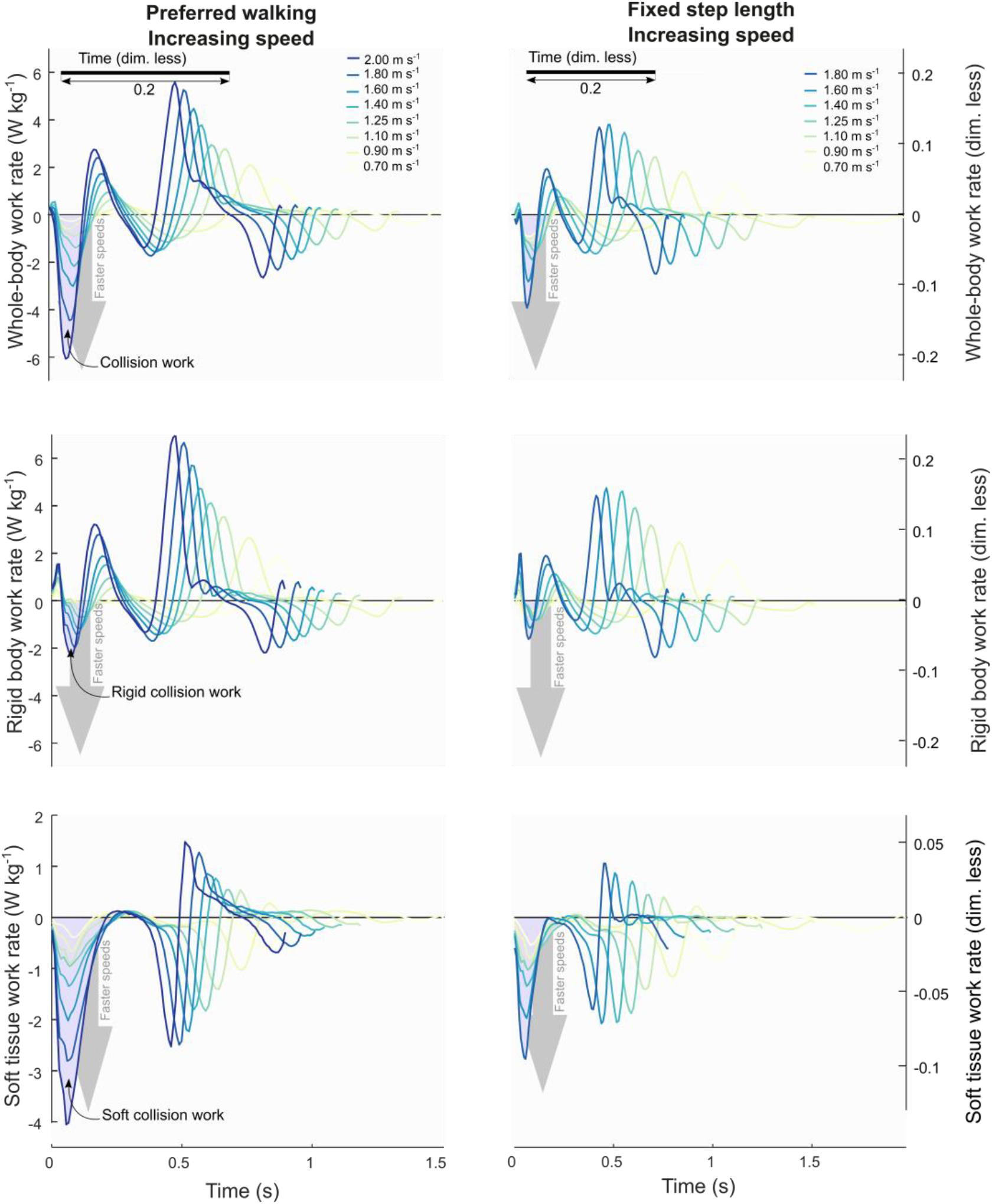
Work rates for walking at increasing speed with (left:) Preferred and (right:) fixed step length. (top to bottom:) Whole-body, Rigid body, and Soft tissue work rates vs. time for one stride, with shaded regions indicating Collision phase. Whole-body work rate is sum of COM and peripheral work rates. Rigid body work is sum of joint work rates from inverse dynamics. Soft tissue work rate is difference between Whole- and rigid body rates.

Negative work increased across conditions according to velocity-based predictions by simple walking model (Fig. 8). As expected, the magnitudes of both Collision work *W*_collision_ (Fig. 8A), soft tissue Collision work *W*_soft,collision_ (Fig. 8B) and full stride negative work *W*_stride_ (Fig. 8C) increased with impact speed 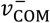 and tangent of COM velocity directional change tan(*δ*_*v*_). All these cases were consistent with the same prediction proportional to 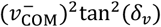 (Eq. 4-6), each with a single coefficient. For example, Collision work *W*_collision_ increased with coefficient *c*_collision_ = 4.512 ± 0.079 (estimate ± 95% confidence interval CI; *p* = 5 · 10^−147^, linear regression, *R*^2^ = 0.81, Fig. 8A). Soft tissue Collision work *W*_soft_ also increased, with a smaller coefficient (*c*_soft_ = 2.849 ± 0.059, *p* = 4 · 10^−130^, linear regression, *R*^2^ = 0.71, Fig. 8B). Full stride negative work *W*_stride_ increased with a larger coefficient (*c*_stride_ = 5.082 ± 0.108, *p* = 1 · 10^−127^, linear regression, *R*^2^ = 0.90, Fig. 8C), and was accompanied by a significant offset in work, (*d*_stride_ = 20.87 ± 0.434 J, mean ± CI).

**Fig. 8:**
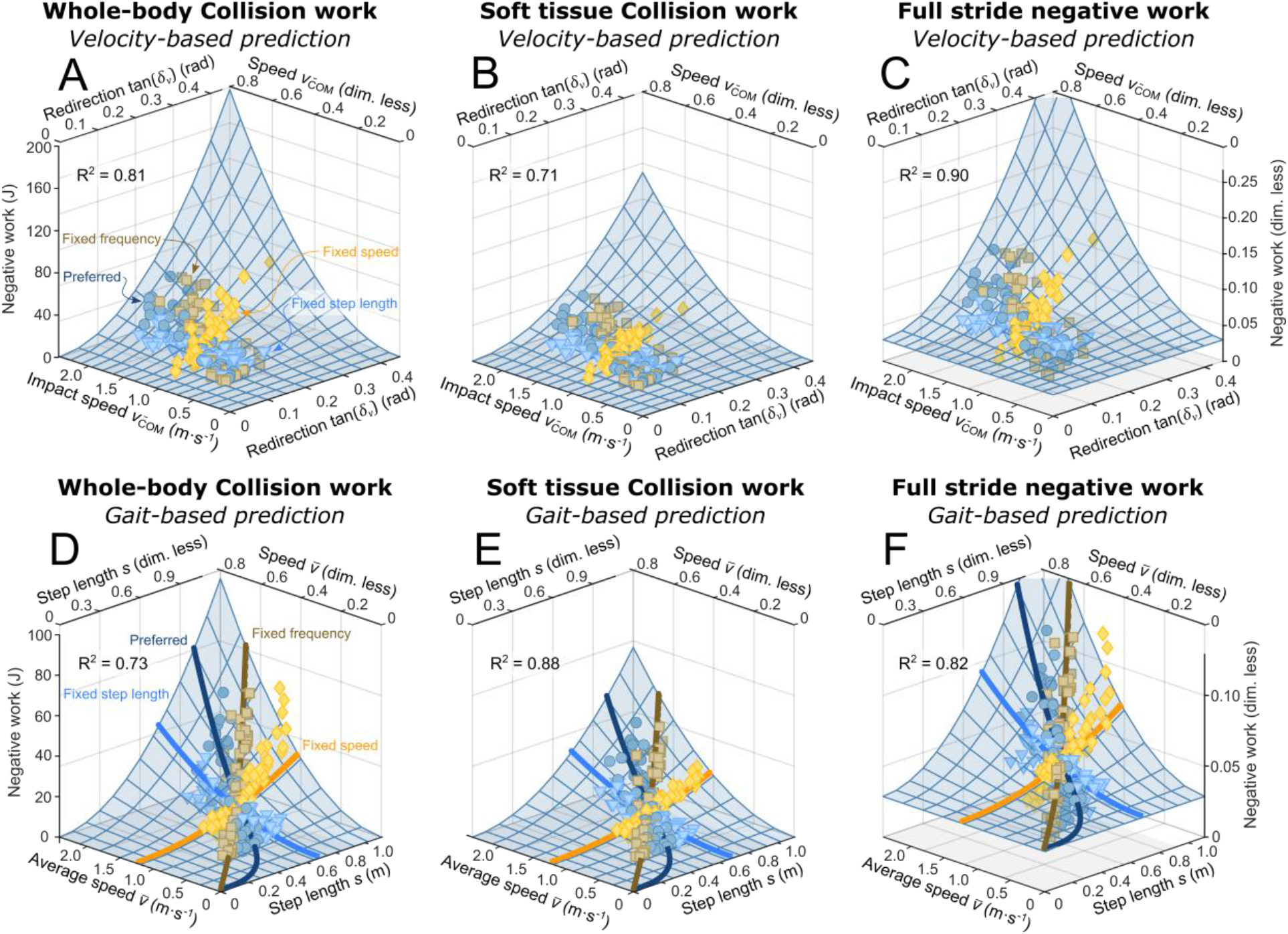
Negative work done by the body and soft tissues during walking, with model prediction (mesh surface). Top row shows Velocity-based predictors, bottom Gait-based predictors. Each row shows Collision work, Soft tissue Collision work, and Full stride negative work. Velocity-based predictors impact speed 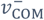 and tangent of COM velocity directional change tan(*δ*_*v*_) for (**A**) whole-body collision (*R*^2^ = 0.82), (**B**) soft tissue collision (*R*^2^ = 0.74) and (**C**) whole-body full stride (*R*^2^ = 0.90). Gait-based predictors average speed 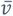 and step length *s* for (**D**) whole-body collision (*R*^2^ = 0.73), (**E**) soft tissue collision (*R*^2^ = 0.88) and (**F**) whole-body full stride (*R*^2^ = 0.82). Each prediction (A-F) was produced with one proportionality coefficient each, and tested with four walking conditions. Step length *s* is normalized by leg length *L*; velocity terms (i.e. 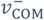 and 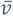) are normalized by 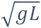, with *g* the gravitational constant.

A comparison of the fitting coefficients reveals that soft tissues accounted for most of the Collision work *W*_collision_, which in turn accounted for most of the full stride negative work *W*_stride_. Quantified by the ratio between coefficients, Collision work *W*_collision_ accounted for 88% of the change in full stride negative work *W*_stride_ (*c*_collision_: *c*_stride_ = 0.88), for the range of walking conditions considered. The main difference was a constant amount of greater full stride negative work *W*_stride_ (20.9 J), not dependent on gait parameters. Similarly, soft tissues accounted for 63% of the Collision work *W*_collision_ (*c*_soft_: *c*_collision_ = 0.63), and 56% of the change in full stride negative work *W*_stride_ (*c*_soft_: *c*_stride_ = 0.56) across conditions considered. Altogether, these results agree with the expectations that soft tissues dissipate substantial energy, mostly during collision phase, which can be quantitatively predicted from impact speed 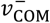 and COM velocity directional change *δ*_*v*_ (Eq. 5).

Negative work increased across conditions according to gait-based predictions by simple walking model (Fig. 8). As expected, the magnitudes of both Collision work *W*_collision_ (Fig. 8D), soft tissue Collision work *W*_soft,collision_ (Fig. 8E), and full stride negative work *W*_stride_ (Fig. 8F) increased with average speed 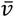 and step length *s*. All these cases were consistent with the same prediction proportional to 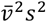 (Eq. 7-9), albeit with different coefficients. For example, Collision work *W*_collision_ increased as described by coefficient *c*′_collision_ = 1.250 ± 0.027 (estimate ± 95% CI; *p* = 5 · 10^−127^, linear regression, *R*^2^ = 0.73, Fig. 8D). Soft tissue Collision work *W*_soft_ also increased, with a smaller coefficient (*c*′_soft_ = 0.828 ± 0.011, *p* = 2 · 10^−178^, linear regression, *R*^2^ = 0.88, Fig. 8E). Full stride negative work *W*_stride_ increased with a larger coefficient (*c*′_stride_ = 1.451 ± 0.042, *p* = 5 · 10^−98^, linear regression, *R*^2^ = 0.82, Fig. 8F), accompanied by an offset (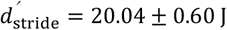, mean ± CI). And as expected for zero net work per stride, full stride *positive* work yielded a comparable coefficient (1.468 ± 0.042), offset (21.89 ± 0.60 J) and overall fit to the same type of proportionality (*R*^2^ = 0.83), meaning that negative and positive work are nearly equal in magnitude. Gait-based coefficients yield predictions for negative work during walking, given subject characteristics (mass *M* and leg length *L*) and gait parameters (speed 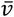 and step length *s*). For example (using Eqn. 8), the predicted amount of negative work done by soft tissues during Collision for the average subject walking at 1.25 m·s^-1^ with a preferred step length of 0.70 m was 6.7 J.

These same fits were also examined on a condition-specific basis, and were found to agree reasonably well with predictions for most conditions (most R^2^ values ≥ 0.5, see Fig. 9). The gait-based predictions (Fig. 8D-F) were evaluated for preferred walking (Fig. 9A), fixed step frequency (Fig. 9B), fixed step length (Fig. 9C), and fixed speed conditions, all using the same single coefficients above (*c*_collision_, *c*_soft,collision_, *c*_stride_). Soft tissue and whole-body negative work matched the predictions best in fixed frequency walking (R^2^ = 0.91-0.95), as expected due to the dominant effect of step length (see Fig. 9B). This was followed by preferred walking (R^2^ = 0.83-0.90), which featured the largest increase in walking speed (see Fig. 9A). Negative work was predicted somewhat less well for fixed speed walking (R^2^ = 0.39-0.66, see Fig. 9D). Soft tissue Collision work *W*_soft,collision_ and full stride negative work *W*_stride_ were reasonably well predicted in fixed step length walking (R^2^ = 0.52-0.68). The fits were relatively poor for whole-body Collision work *W*_collision_ at fixed step lengths (R^2^ = 0.13, see Fig. 9C), but this was because work changed little across this condition (9.9 J at most), and not because of substantial absolute error in the fit. Separate from these predictions, the Rigid body Collision work was small in all conditions, taking up only 28% of (whole-body) Collision work *W*_collision_ (averaged across all conditions). Altogether, data agreed with predictions mainly for the preferred, fixed frequency, and fixed speed predictions, where there was generally more change in dissipation across trials.

**Fig. 9:**
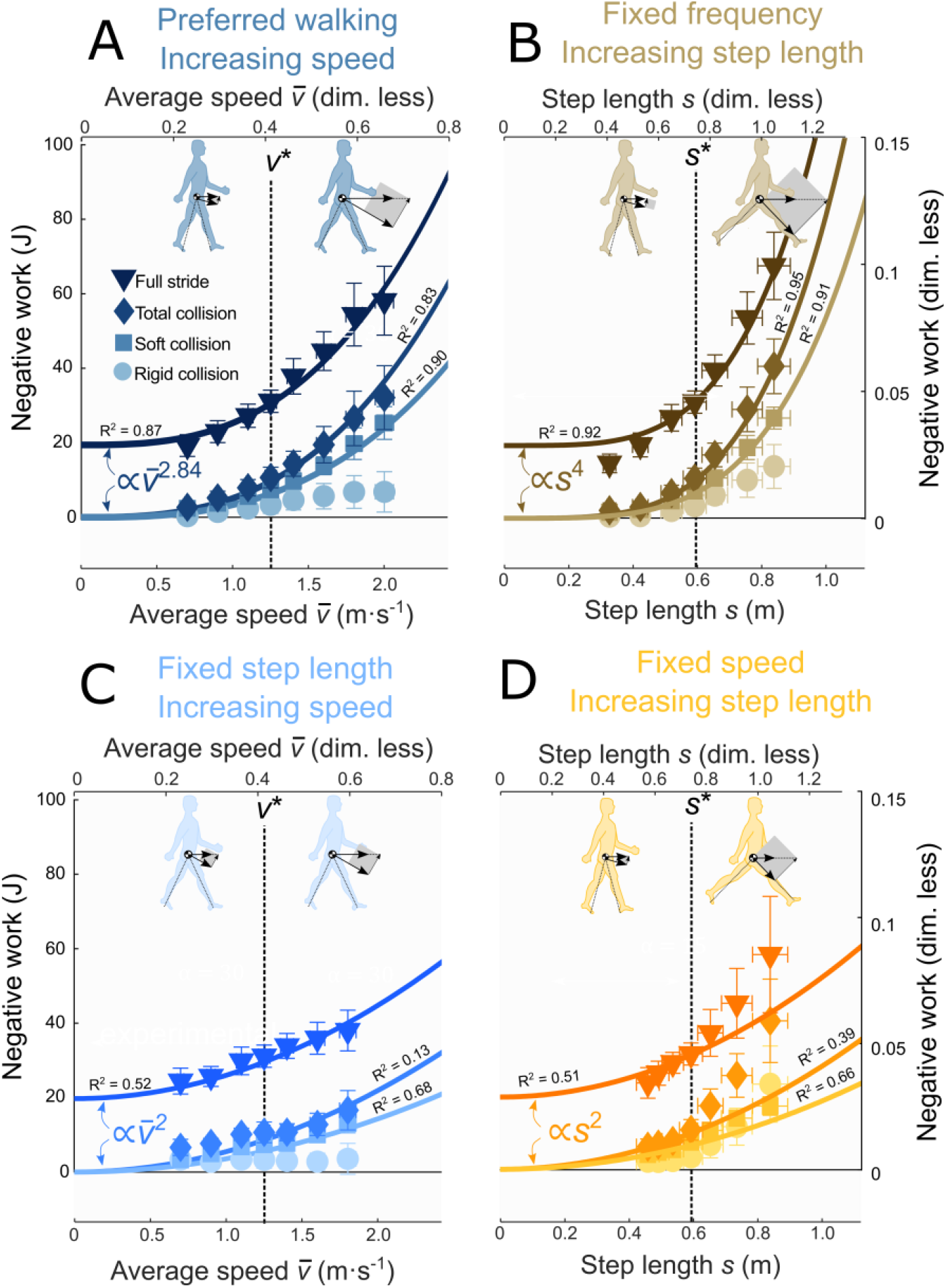
Condition-specific effects of step length and speed on negative work during walking. A: Negative work versus average speed for preferred walking (R^2^ = 0.83-0.90). B: Negative work versus step length for fixed frequency walking (R^2^ = 0.91-0.95). C: Negative work versus average speed for fixed step length walking (R^2^ =0.13–0.68). D: Negative work versus step length for fixed speed walking, with poorer fit to predictions (R^2^ = 0.39-0.66). All four conditions were tested against a single model with one proportionality coefficient (Fig. 8D-F) for each of three quantities: Full stride negative work, total negative Collision work, and soft tissue Collision work (*c*_stride_, *c*_collision_, *c*_soft,collision_). Step length *s* is normalized by leg length *L*; average speed 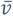 is normalized by 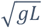, with *g* the gravitational constant.

## Discussion

The current study aimed at testing whether negative work by the whole body or passive soft tissues varies systematically at various combinations of walking speed and step length. We tested whether the negative work done by soft tissues during Collision (“soft tissue Collision work”), and the whole-body negative work over a full stride, were both proportional to whole-body Collision work. And in turn, we also tested whether Collision work would increase as predicted by a simple model, with step length squared multiplied by walking speed squared. These quantities were found to agree reasonably well with the model. We next examine the results considering potential underlying mechanisms, as well as the implications for biomechanical analysis of human locomotion.

We found that soft tissues do substantial amounts of negative work over a wide range of walking conditions. Soft tissue work has previously been related to walking speed during preferred walking (Zelik & Kuo, 2010), but not for other conditions, and not relative to negative work done by the whole body. We here show that soft tissues account for most (about 65%) of the negative work done by the whole-body during Collision, over a variety of conditions quite different from preferred walking (Fig. 8). The negative Collision work not performed by soft tissues may be performed by a combination of active dissipation by muscle, and tendon passively storing some energy elastically, and perhaps returning during Rebound (e.g., Fig. 4C). Our data are insufficient to quantify the actual amount of elastic return, which has suggested to be quite substantial and attributable to the knee (Shamaei et al., 2013). But soft tissues appear to be well damped, with little indication of elastic return (see Fig. 6-7). If soft tissue Collision losses (about 6.7 J at typical 1.25 m · s^−1^) were restored by active positive work at 25% efficiency (Margaria, 1968), it could account for about 31% of the net metabolic power of about 2.3 W · kg^−1^ at that speed (Kuo et al., 2005). Soft tissue work is therefore an important dissipative contributor to negative work, and ultimately to the metabolic cost of walking.

We also found that the negative work of the entire stride is related to the Collision. Both quantities (*W*_stride_ and *W*_collision_) increased proportionately, with approximately the same power law with respect to either velocity- or gait-based predictors (Fig. 8). The Collision accounted for about 88%, and soft tissues for about 57%, of the changes in negative work over an entire stride. This leaves a relatively small, 12% contribution from other phases to changes in overall negative work, albeit still in proportion to Collision. There was also a substantial offset in the full stride negative work, accounting for as much as 87% of the total negative work at low speeds (below nominal). We interpret the offset as arising from other factors not considered here, such as the contribution of step width to Collision (Donelan et al., 2001, 2002), and motion of the swing leg (Doke et al., 2005) and other parts of the body. The negative work of Pre-load appears to be associated with elastic loading of the Achilles tendon, prior to subsequent release as part of Push-off (Fukunaga et al., 2001; Zelik et al., 2014). That loading appears driven in part by the dynamical interactions triggered by Collision, as is the overall positive work of the body over the full stride.

These changes are mechanistically predictable over a wide range of walking conditions. The mechanistic stimulus is the vector velocity of the COM, which is redirected during the step-to-step transition. Negative Collision work was predicted from velocity’s magnitude 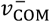 and directional change *δ*_*v*_ (Eq. 4), which both varied substantially across walking conditions. Despite a greater than two-fold variation in each of the gait parameters, the Collision work was predicted reasonably well by a single empirical coefficient *c*_soft_ (*R*^2^ = 0.81), and similarly for soft tissue Collision work and whole-body negative over the stride. A drawback is that such predictions require the pre-impact COM velocity 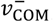, and so we also tested more convenient gait-based variables, namely speed 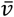 and step length *s*, that are more typically known or specified prior to experiment. The gait-based predictions rely on assumptions such as the small-angle approximation, and that impact velocity is proportional to average speed. However, we found gait-based predictions to fit data about as well as velocity-based predictions (R^2^ = 0.88 vs. R^2^ = 0.74). The exception was for large step lengths during fixed speed walking, which feature substantial directional changes *δ*_*v*_ but relatively less soft tissue Collision work (see Fig. 9). Humans appear to behave less like inverted pendulums when walking with atypically high step lengths and low step frequencies. Although the fits for this condition are not as good as for some other conditions (with R^2^ between 0.39 and 0.66), this is a consequence of performing a single fit for all conditions, some of which entailed much more work and therefore had greater influence on the proportionality coefficient. Better fits could be obtained with a separate coefficient for each specific condition, but our aim was to test a single model across all conditions. With the single coefficient limitation in mind, both soft tissue dissipation and whole-body negative work can be predicted reasonably from gait parameters for walking speed and step length, particularly for conditions similar to normal preferred walking.

These predictions arise from a simple walking model that predicts the work needed to redirect the COM velocity during the step-to-step transition. It predicts general trends arising from pendulum-like walking (Eqns 1-3), and not absolute work quantities, which required empirical proportionality coefficients. However, only a single proportionality coefficient could reasonably predict the work for all four different walking conditions, whether for soft tissues, overall collision, or full stride (e.g.,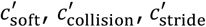). The model prediction does not distinguish between active and passive work, but we found that soft tissue dissipation was proportional to whole-body negative work. It appears that humans are quite systematic in distributing work between passive soft tissues and active muscle-tendons. Thus, the combination of a simple model and only a few empirical coefficients, unites the effects of a variety of gait parameters on negative work. At the same time, the simple model applies less well to the fixed-speed combinations of long step lengths and slow step frequencies discussed above. But for step lengths and frequencies at or near preferred conditions, the inverted pendulum model explains the negative work of soft tissues and the whole body well, during both collision and the full stride.

These findings also help to reveal that traditional inverse dynamics analysis is least accurate during Collision. Rigid body work accounted for only about 28% of total negative work following the impact at heel strike, across a wide range of walking conditions. The ratio of rigid body negative work to total negative work during Collision was largest during fixed speed walking (36%) and smallest during preferred walking (21%). This is also corroborated by observed net rigid body work for a full stride not being zero as expected for periodic gait, but rather positive (Fig. 5) and increasing with greater speeds or step lengths. This can largely be explained by the Collision phase, when soft tissues are most dissipative, and which rigid body work cannot capture (compare Figs. 4A and B). Rigid body work appears to be improved by the use of 6D inverse dynamics, which can in principle capture some of the work performed between neighboring segments or within the joints (Honert & Zelik, 2019). However, inverse dynamics seems quite inaccurate for quantifying the work performed during Collision, which occurs within the first 200 ms or so after heel strike. The present study indicates the specific gait conditions and amount of work (Eqs. 7 – 9, each with an empirical coefficient) not quantified by inverse dynamics.

There are also limitations to the quantification and interpretation of soft tissue dissipation. For example, we observed a large negative burst of soft tissue work followed by a large positive burst during pre-load and push-off respectively (Figs. 6 and 7). Soft tissues cannot actively perform positive work, and so it is possible that the negative-positive sequence is from passive, elastic energy return by soft tissues, first compressed and stored during pre-load and then released during push-off, in unknown amount. The timing and magnitude of positive work seem consistent with that interpretation, but another possibility is that the work is not truly caused by soft tissues, but by unmodeled rigid body joints. In our analysis, we estimate soft tissue dissipation from the energy not accounted for by the rigid body model. For example, we did not measure the metatarsophalangeal (MTP) joint, which may store and return energy during walking (Farris et al., 2019), and could potentially be included in a multi-segment (Farris et al., 2019) or deformable foot model (e.g. Takahashi et al., 2012) compatible with inverse dynamics. There remains the question whether such action is actively performed by muscle or passively by tendons, which may be addressed through techniques such as ultrasound imaging (Fukunaga et al., 2001). Thus, interpretation of soft tissue work can also depend on rigid body assumptions and on passive elasticity.

Soft tissue dissipation might superficially seem preferable to avoid. All negative work, whether by soft tissue dissipation or by active muscle, needs to be restored by an equal amount of positive work (Riddick & Kuo, 2020) in steady level locomotion, at the cost of metabolic energy. However, negative work is necessary during the step-to-step transition to redirect the COM velocity. Doing this necessary negative work with soft tissues instead of active muscles may be more economical, as muscles require metabolic energy even for negative work (Abbott et al., 1952). The possible economy of soft tissue dissipation is supported by the lower mass-normalized metabolic cost of walking for obese than healthy individuals (Fu et al., 2015). Soft tissues also enable a softer impact with the ground (Pain & Challis, 2006), which may help in avoiding damage or injury to other tissues. For example, high knee adduction moment impulse is considered a risk factor for knee osteoarthritis (Bennell et al., 2011), whereas high vertical loading rate is considered a risk factor for tibial stress syndrome (Milner et al., 2006). The human nervous system appears to apportion some negative work to soft tissues, and some to muscle-tendons under active control. For example, humans prefer a jump landing that requires 37% more muscle-tendon dissipation than minimally necessary (Zelik & Kuo, 2012). The amount of soft tissue dissipation may also have other effects such as on the stability of walking (Masters & Challis, 2020). The distribution between active and passive dissipation may therefore be relevant to metabolic cost and a variety of additional mechanical effects.

## Conclusion

Soft tissue dissipation during walking accounts for 56% of the variation in total negative work during walking. Both soft tissue and total negative work increase in consistent relative proportion, and with the square of walking speed and step length as predicted by a simple model of pendulum-like walking. The model mechanistically explains how negative work is necessary to redirect the body’s velocity between pendulum-like steps. Across a variety of conditions, experimental data reveal substantial soft tissue dissipation during walking, in predictable amount not captured by rigid body inverse dynamics analysis. In steady gait, negative and positive work are performed in equal magnitude, so that dissipative soft tissue work also requires active positive work that costs metabolic energy.

## Acknowledgements

The authors thank Emily Mundinger for contributions to data analysis.

## Data availability

The entire data set will become available at a later stage as part of another paper.

## References

Abbott, B. C., Bigland, B., & Ritchie, J. M. (1952). The physiological cost of negative work. The Journal of Physiology, 117(3), 380–390.

Adamczyk, P. G., Collins, S. H., & Kuo, A. D. (2006). The advantages of a rolling foot in human walking. Journal of Experimental Biology, 209, 3953–3963.

Adamczyk, P. G., & Kuo, A. D. (2009). Redirection of center-of-mass velocity during the step-to-step transition of human walking. Journal of Experimental Biology, 212, 2668–2678.

Baines, P. M., Schwab, A. L., & van Soest, A. J. (2018). Experimental estimation of energy absorption during heel strike in human barefoot walking. PLoS ONE, 13(6). https://doi.org/10.1371/journal.pone.0197428

Bennell, K. L., Bowles, K.-A., Wang, Y., Cicuttini, F., Davies-Tuck, M., & Hinman, R. S. (2011). Higher dynamic medial knee load predicts greater cartilage loss over 12 months in medial knee osteoarthritis. Annals of the Rheumatic Diseases, 70(10), 1770–1774. https://doi.org/10.1136/ard.2010.147082

Davis, R. B., Õunpuu, S., Tyburski, D., & Gage, J. R. (1991). A gait analysis data collection and reduction technique. Human Movement Science, 10(5), 575–587. https://doi.org/10.1016/0167-9457(91)90046-Z

Doke, J., Donelan, J. M., & Kuo, A. D. (2005). Mechanics and energetics of swinging the human leg. The Journal of Experimental Biology, 208(Pt 3), 439–445. https://doi.org/10.1242/jeb.01408

Donelan, J. M., Kram, R., & Kuo, A. D. (2001). Mechanical and metabolic determinants of the preferred step width in human walking. Proceedings. Biological Sciences / The Royal Society, 268(1480), 1985–1992. https://doi.org/10.1098/rspb.2001.1761

Donelan, J. M., Kram, R., & Kuo, A. D. (2002). Mechanical work for step-to-step transitions is a major determinant of the metabolic cost of human walking. Journal of Experimental Biology, 205(23), 3717–3727.

Farris, D. J., Kelly, L. A., Cresswell, A. G., & Lichtwark, G. A. (2019). The functional importance of human foot muscles for bipedal locomotion. Proceedings of the National Academy of Sciences, 116(5), 1645–1650. https://doi.org/10.1073/pnas.1812820116

Fu, X.-Y., Zelik, K. E., Board, W. J., Browning, R. C., & Kuo, A. D. (2015). Soft Tissue Deformations Contribute to the Mechanics of Walking in Obese Adults. Medicine and Science in Sports and Exercise, 47(7), 1435–1443. https://doi.org/10.1249/MSS.0000000000000554

Fukunaga, T., Kubo, K., Kawakami, Y., Fukashiro, S., Kanehisa, H., & Maganaris, C. N. (2001). In vivo behaviour of human muscle tendon during walking. Proceedings of the Royal Society B— Biological Sciences, 268(1464), 229–233. https://doi.org/10.1098/rspb.2000.1361

Geyer, H., Seyfarth, A., & Blickhan, R. (2006). Compliant leg behaviour explains basic dynamics of walking and running. Proc Biol Sci, 273, 2861–2867.

Grieve, D. W. (1968). Gait patterns and the speed of walking. Biomedical Engineering, 3, 119–122.

Hayes, W. C., & Mockros, L. F. (1971). Viscoelastic properties of human articular cartilage. J Appl Physiol, 31, 562–568.

Honert, E. C., & Zelik, K. E. (2019). Foot and shoe responsible for majority of soft tissue work in early stance of walking. Human Movement Science, 64, 191–202. https://doi.org/10.1016/j.humov.2019.01.008

Kuo, A. D. (2001). A simple model of bipedal walking predicts the preferred speed-step length relationship. Journal of Biomechanical Engineering, 123(3), 264–269.

Kuo, A. D., Donelan, J. M., & Ruina, A. (2005). Energetic consequences of walking like an inverted pendulum: Step-to-step transitions. Exercise and Sport Sciences Reviews, 33(2), 88.

Margaria, R. (1968). Positive and negative work performances and their efficiencies in human locomotion. Internationale Zeitschrift Für Angewandte Physiologie, Einschliesslich Arbeitsphysiologie, 25(4), 339–351.

Masters, S., & Challis, J. (2020). Soft tissue vibration: A biologically-inspired mechanism for stabilizing bipedal locomotion. Bioinspiration & Biomimetics. https://doi.org/10.1088/1748-3190/abd624

Milner, C. E., Ferber, R., Pollard, C. D., Hamill, J., & Davis, I. S. (2006). Biomechanical factors associated with tibial stress fracture in female runners. Medicine and Science in Sports and Exercise, 38(2), 323–328. https://doi.org/10.1249/01.mss.0000183477.75808.92

Minetti, A. E., & Belli, G. (1994). A model for the estimation of visceral mass displacement in periodic movements. J Biomech, 27, 97–101.

Pain, M. T., & Challis, J. H. (2001). The role of the heel pad and shank soft tissue during impacts: A further resolution of a paradox. J Biomech, 34, 327–333.

Pain, M. T., & Challis, J. H. (2006). The influence of soft tissue movement on ground reaction forces, joint torques and joint reaction forces in drop landings. J Biomech, 39, 119–124.

Riddick, R. C., & Kuo, A. D. (2020). Mechanical work accounts for most of the energetic cost in human running. Scientific Reports.

Schmitt, S., & Günther, M. (2011). Human leg impact: Energy dissipation of wobbling masses. Archive of Applied Mechanics, 81(7), 887–897. https://doi.org/10.1007/s00419-010-0458-z

Shamaei, K., Sawicki, G. S., & Dollar, A. M. (2013). Estimation of quasi-stiffness of the human knee in the stance phase of walking. PloS One, 8(3), e59993. https://doi.org/10.1371/journal.pone.0059993

Takahashi, K. Z., Kepple, T. M., & Stanhope, S. J. (2012). A unified deformable (UD) segment model for quantifying total power of anatomical and prosthetic below-knee structures during stance in gait. Journal of Biomechanics, 45(15), 2662–2667. https://doi.org/10.1016/j.jbiomech.2012.08.017

Vaughan, C., Davis, B., & Jeremy, C. (1992). Dynamics of human gait. Human Kinetics Publishers Champaign, Illinois.

Virgin, W. J. (1951). Experimental investigations into the physical properties of the intervertebral disc. J Bone Joint Surg Br, 33-B, 607–611.

Zelik, K. E., Huang, T.-W. P., Adamczyk, P. G., & Kuo, A. D. (2014). The role of series ankle elasticity in bipedal walking. Journal of Theoretical Biology, 346, 75–85. https://doi.org/10.1016/j.jtbi.2013.12.014

Zelik, K. E., & Kuo, A. D. (2010). Human walking isn’t all hard work: Evidence of soft tissue contributions to energy dissipation and return. Journal of Experimental Biology, 213(24), 4257–4264. https://doi.org/10.1242/jeb.044297

Zelik, K. E., & Kuo, A. D. (2012). Mechanical work as an indirect measure of subjective costs influencing human movement. PloS One, 7(2), e31143. https://doi.org/10.1371/journal.pone.0031143

Zelik, K. E., Takahashi, K. Z., & Sawicki, G. S. (2015). Six degree-of-freedom analysis of hip, knee, ankle and foot provides updated understanding of biomechanical work during human walking. The Journal of Experimental Biology, 218(Pt 6), 876–886. https://doi.org/10.1242/jeb.115451

